# Intra- and interpatient evolution of enterovirus D68 analyzed by whole-genome deep sequencing

**DOI:** 10.1101/420836

**Authors:** Robert Dyrdak, Monika Mastafa, Emma B. Hodcroft, Richard A. Neher, Jan Albert

**Affiliations:** Department of Clinical Microbiology, Karolinska University Hospital, Stockholm, Sweden; Department of Microbiology, Tumor and Cell Biology, Karolinska Institute, Stockholm, Sweden; Biozentrum, University of Basel, Basel, Switzerland; Swiss Institute of Bioinformatics, Basel, Switzerland

## Abstract

Worldwide outbreaks of enterovirus D68 (EV-D68) in 2014 and 2016 have caused serious respiratory and neurological disease. To investigate diversity, spread, and evolution of EV-D68 we performed near full-length deep sequencing in 54 samples obtained in Sweden during the 2014 and 2016 outbreaks. In most samples, intrapatient variability was low and dominated by rare synonymous variants, but three patients showed evidence of dual infections with distinct EV-D68 variants from the same subclade. Interpatient evolution showed a very strong temporal signal, with an evolutionary rate of 0.0039 ± 0.0001 substitutions per site and year. Phylogenetic trees reconstructed from the sequences suggest that EV-D68 was introduced into Stockholm several times during the 2016 outbreak. Putative neutralization targets in the BC and DE loops of the VP1 protein were slightly more diverse within-host and tended to undergo more frequent substitution than other genomic regions. However, evolution in these loops did not appear to have been driven the emergence of the 2016 B3-subclade directly from the 2014 B1-subclade. Instead, the most recent ancestor of both clades was dated to 2009. The study provides a comprehensive description of the intra- and interpatient evolution of EV-D68, including the first report of intrapatient diversity and dual infections. The new data along with publicly available EV-D68 sequences are included in an interactive phylodynamic analysis on nextstrain.org/enterovirus/d68 to facilitate timely EV-D68 tracking in the future.

## INTRODUCTION

Enterovirus D68 (EV-D68) has been recognized as an emerging pathogen after recent worldwide outbreaks of serious respiratory and neurological disease. The virus was discovered in 1962 in California, USA in four children with pneumonia (Schieble *et al*., 1967). Prior to 2014, EV-D68 was reported only sporadically with a total of 699 confirmed cases in Europe, Africa, and southeast Asia between 1970 to 2013 (as reviewed by Holm-Hansen *et al*. (2016)). But in 2014, 2,287 of cases with EV-D68 infection were reported, mainly in North America and Europe, but also in southeast Asia and Chile (Holm-Hansen *et al*., 2016). Following the 2014 outbreak, another wave of EV-D68 infections was observed in 2016 with reports of outbreaks in several parts of the world, including Europe (Barnadas *et al*., 2017; Dyrdak *et al*., 2016; Knoester *et al*., 2017; Piralla *et al*., 2018), USA (Messacar *et al*., 2017; Wang *et al*., 2017), Argentina (Ruggieri *et al*., 2017), and Taiwan (Wei *et al*., 2018).

Even though EV-D68 was rarely reported before the recent outbreaks, a high prevalence of neutralizing antibodies have reported in samples collected before 2014 (Vogt and Crowe Jr, 2018). This indicates that infection with EV-D68 has been common also before the outbreaks in 2014 and 2016.

The main clinical presentation of EV-D68 infection is mild to severe respiratory symptoms (Holm-Hansen *et al*., 2016), but in rare cases EV-D68 can also cause a poliomyelitis-like disease termed acute flaccid myelitis (AFM) (Dyda *et al*., 2018; Messacar *et al*., 2018). An increase of this otherwise unusual clinical presentation coincided with the outbreak of EV-D68 in 2014. In animal studies, strains from 2014 were more virulent than the prototype Fermon strain isolated in 1969 (Hixon *et al*., 2017b; Zhang *et al*., 2018a).

EV-D68, as other enteroviruses, has a genome size of approximately 7.5 kb and codes for a polyprotein, which is processed into four structural (VP1-4), and seven non-structural proteins (2A-C, 3A-D). Phylogenetically EV-D68 is divided into three clades (A, B and C) (Tokarz *et al*., 2012), and the B clade is further subdivided into three sub-clades or lineages (B1, B2 and B3) (Gong *et al*., 2016). The 2014 and 2016 outbreaks were caused by viruses belonging to the B1 and B3 subclades, respectively (Barnadas *et al*., 2017; Dyrdak *et al*., 2016; Knoester *et al*., 2017; Piralla *et al*., 2018; Wang *et al*., 2017; Wei *et al*., 2018; Zhang *et al*., 2016). Similar to other RNA viruses, EV-D68 has been reported to have a high substitution rate with 6.2·10^−3^ (Tokarz *et al*., 2012) or 5.12 · 10^−3^ (Ny *et al*., 2017) substitutions per site per year in the highly variable VP1 gene.

Mechanisms of immune protection and immune escape for EV-D68 are not well understood. Neutralizing antibodies have been shown to be protective in animal models (Dai *et al*., 2018; Hixon *et al*., 2017a,b; Zhang *et al*., 2018a,b). Based on homology with the neutralizing sites NIm-IA and NIm-IB on human rhinovirus 14 (HRV-14), regions on the BC and DE loops in EV-D68 have been proposed as putative epitopes for neutralizing antibodies (Liu *et al*., 2015). However, also other sites in VP1 and other proteins may be of importance. Thus, amino acid variability in regions flanking the VP1 loops were reported to correlate with differences in neutralisation sensitivity between the prototype Fermon strain and strains from 2014 (Zhang *et al*., 2015). For coxsackie virus B4, it was shown that a deletion in the BC loop reduced the neutralizing effect of CVB4-specific antisera (McPhee *et al*., 1994). VP2 and VP3 may also contain neutralizing epitopes as demonstrated in early poliovirus studies (Minor *et al*., 1986) and as recently reviewed for the enterovirus A genus (Fang and Liu, 2018). Little is known about the contribution of cellular immunity to the clearance and evolution of EV-D68 (and other enteroviruses).

Here, we report deep sequencing of near full-length genomes on the Illumina next-generation sequencing (NGS) platform and analyze intra- and interpatient evolution of EV-D68 during the recent outbreaks. We show that within-host diversity is typically low, but that dual infection is not uncommon, suggesting high infection incidence during the outbreaks. To facilitate exploration of the data and future analyses, we combined the whole genome sequences reported here with publicly available genomes and implemented an interactive visualization in `nextstrain` available at nextstrain.org/enterovirus/d68.

## MATERIALS & METHODS

### Study population and samples

For this study we used 54 nasopharyngeal and 5 other respiratory samples from two earlier studies on EV-D68 outbreaks in Stockholm in 2014 and 2016 (Dyrdak *et al*., 2016, 2015). The 59 samples were drawn from 54 patients. Six patients had been sampled in 2014, and 48 patients (53 samples) in 2016. Thirty of the patients were female. The median age was 3.6 years (range 2 months – 63 years). Five patients were sampled twice during their acute illness (0, 1, 1, 2 and 7 days apart).

All, but one, samples were submitted from hospitals in Stockholm. However, we cannot exclude the possibility that a few samples might be from patients who had been transferred for care to Stockholm from other parts of Sweden.

Details about the patient demographics are given in supplementary table S2.

The collection dates of the samples from the outbreak in 2016 ranged between August 19 to September 13. The samples had tested positive for EV-D68 at the Department of Clinical Microbiology at Karolinska University Hospital, by EV-D68 specific realtime PCR and/or partial VP4/VP2 sequencing. For details see (Dyrdak *et al*., 2016, 2015). A few EV-D68-positive samples from the earlier studies were not included in this study because no sample material remained or because the virus levels were very low as evaluated by the cycle threshold (Ct) values in the EV-D68 realtime PCR. The Ct values for the included samples ranged from 13.9 to 33.9 (median 22.3). Details about the study samples are given in supplementary table S3.

The study was reviewed and approved by the Regional Ethical Review Board in Stockholm, Sweden (registration no. 2017/1317–32).

### Near full-length EV-D68 amplification

Primers were designed to cover almost the entire EV-D68 genome in four overlapping fragments (F1-F4) that were approximately 2,000 base pairs long (Table S1). An alignment of published EV-D68 genomes was used to design forward and reverse primers targeting highly conserved regions of the EV-D68 genome, with similar melting temperatures, and with minimal tendency for hairpin and primer-dimer formation. Inner nested primers for fragment F1 were designed for template quantification.

RNA was manually extracted using the RNeasy Lipid Tissue Mini Kit (Qiagen Cat. No. 74804). For each nasopharyngeal sample, 200 *μ*l was taken and mixed with 1 ml Qiazol, 200 *μ*l chloroform and 5 *μ*l carrier-RNA. After centrifugation and several washing steps, each aliquot was eluted with 100 *μ*l RNase-free water and were either stored at −70°C or used directly for PCR amplification.

Each sample was amplified in duplicate by one-step RT-PCR for each of the four overlapping fragments. The RT-PCR mixture contained: 11 *μ*l of RNA template, 0.5 *μ*M of forward and reverse PCR primer, 1 ng/*μ*l random hexamers, 1 *μ*l Superscript III RT/Platinum Taq HiFi Enzyme mix (Invitrogen, Stockholm, Sweden), 25 *μ*l 2x reaction mix, and RNase-free water to a total volume of 50 *μ*l. The PCR cycling profile was: ×1: 30 min at 50°C, 2 min at 94°C (cDNA synthesis); ×30: 15 sec at 94°C, 30 sec at 50°C, 90 sec at 68°C (PCR-amplification); ×1: 5 min at 68°C, ∞ at 4°C (final extension).

To approximately quantify the number of input EV-D68 RNA templates for sequencing, a dilution series of the RNA (1:10, 1:100, 1:1000) was amplified in duplicate with nested primers for fragment F1. EV-D68 RNA templates were enumerated by the Poisson distribution formula. The outer PCR is described above. The nested PCR mixture contained: 2.5 *μ*l of product from the outer PCR, 0.2 *μ*M of forward and reverse PCR primer, 1 unit of Platinum Taq HiFi (Invitrogen, Stockholm, Sweden), 5 *μ*l 10x reaction mix, and RNase-free water to a total volume of 50 *μ*l. The cycling profile was: ×1: 2 min at 94°C, (denaturation); ×30: 15 sec at 94°C, 30 sec at 50°C, 90 sec at 68°C (PCR-amplification); ×1: 6 min at 68°C, at 6 min (final extension).

Duplicates of each fragment were pooled and purified with AGENCOURT AMPure XP PCR purification kit and quantified with Qubit assays (Q32851, Life Technologies). Purified DNA from each fragment were diluted to the same concentration, pooled and sent to the Clinical Genomics Unit at Science for Life Laboratory (SciLife-Lab, Stockholm, Sweden) for sequencing.

### DNA library preparation and sequencing

The DNA preparations were quantified using Quant-iTTM dsDNA High-Sensitivity Assay Kit and Tecan Spark 10M plate reader. 1 *μ*l of DNA (approximately 0.5-2.0 ng/*μ*l) was used in the tagmentation reaction using Nextera chemistry (Illumina) to yield fragments larger than 150 bp. The tagmented library underwent 11 cycles of PCR with single-end indexed primers (IDT Technologies) followed by purification using Seramag beads according to the protocol. The library was quantified using Quant-iTTM dsDNA High-Sensitivity Assay Kit and Tecan Spark 10M plate reader and pair-end (2×101 bp) sequenced to a depth of 100,000 – 1,000,000 reads per sample on a HiSeq 2500 Illumina sequencer. Base calling and demultiplexing was done using bcl2fastq v1.87, without allowing any mismatch in the index sequence.

### Quality controls experiments

Quality controls were prepared from two samples with low Ct values (SWE_027_160829 and SWE_037_160829, having Ct values 14.62 and 19.27, respectively). The samples were diluted 1:100 in a pool of virus-negative nasopharyngeal samples and stored at −70° C in 200 *μ*l aliquots. The enterovirus RNA concentration of the samples was quantified by limiting dilution in 10 replicas of ten-fold dilutions, from 1:10 to 1:100,000 with an inhouse enterovirus realtime-PCR (Tiveljung-Lindell *et al*., 2009). Based on Poisson calculations, the samples were estimated to have 4,400 and 240 EV-D68 RNA copies per *μ*l, respectively, prior to dilution. The two samples were RNA extracted, PCR amplified and sequenced twice in two separate runs. The number of EV-D68 RNA templates in each PCR reaction was approximately 48,400 and 2,640, respectively.

### Read filtering, mapping and analysis

Sequencing reads were trimmed using TrimGalore (Krueger, 2015; Martin, 2011) and mapped against the sequence `KX675261.1` using `bwa mem` (Li and Durbin, 2009). We used the `pysam` wrapper to samtools to generate a pile-up of all reads and quantify insertions relative to the reference. Custom python scripts were used to generate consensus sequences and quantify intrasample diversity from the pile-ups. All these steps were chained using the workflow engine `snakemake` (Köster and Rahmann, 2012). The entire pipeline is available github.com/neherlab/EV-D68_analysis_Dyrdak_2019 and uses a code from a separate project github.com/neherlab/SVVC. Commit `81068ee` was used to analyze the data along with commit `a2cbb35` of `SVVC`. To analyze linkage disequilibrium, we calculated genotype frequencies *p*_12_ at pairs of positions that were covered by a read pair more than 100-fold and calculated their correlation *D*_12_ = *p*_12_ – *p*_1_*p*_2_ (aka linkage disequilibrium). This correlation was normalized such that maximal linkage corresponds to *D*_12_ = 1 and complete repulsion to *D*_12_ = −1.

The consensus sequences for successfully sequenced samples have been deposited in GenBank (accession numbers MH674111–MH674166 and MH844544).

The scripts that analyze diversity in pile-ups and generate most of the the figures of this study are available at github.com/neherlab/EV-D68_analysis_Dyrdak_2019.

### Phylodynamics analysis

Sequences generated in this study were combined with sequences and metadata of all EV-D68 genomes with a length of at least 6,000 bp available on Virus Pathogen Resource (ViPR) (Pickett *et al*., 2011) as of 2018-09-02. The combined data set was analyzed using `nextstrain` (Hadfield *et al*., 2018) and snakemake (Köster and Rahmann, 2012). The `augur` pipeline was run using the aligner MAFFT (Katoh *et al*., 2002), the tree-builder (Nguyen *et al*., 2015), and the phylodynamic package TreeTime (Sagulenko *et al*., 2018). The repository detailing the analysis pipeline is available at https://github.com/neherlab/enterovirus_nextstrain. Sub-genogrouping of sequences was done using the Enterovirus Genotyping Tool Version 1.0 (Kroneman *et al*., 2011). The resulting analysis is visualized with auspice and is available at nextstrain.org/enterovirus/d68.

Five sequences were excluded from the phylodynamic analysis because they were deemed problematic. The following three sequences are within a few mutations from other sequences sampled several years earlier or later: EV-D68/Homoj3apiens/USA/U797/2007 is identical to strain EV-D68/Homo_sapiens/USA/C7791/2014; EV_D68/Homo_sapiens/USA/0622a/2012 is very similar to strains sampled in 2003; EV_D68/Homo_sapiens/USA/M078/2009 is very similar to strains sampled in 2014 in the USA. Given the high evolutionary rate of enteroviruses, such stasis over many years is very unlikely (we expect about 30 changes per year) and these sequences are likely dated incorrectly. The case CA/RESP/10_786 is less clear, but the sequence is several standard deviations less diverged from the root than expected given its collection date. USA/TX/2014_19267 is likely a recombinant sequence.

Mixed effects model of evolution (MEME) analysis (Murrell *et al*., 2012) using the DataMonkey package (Weaver *et al*., 2018) was used to analyze for episodic diversifying selection.

## RESULTS

Fifty-four of 59 samples yielded a coverage of at least 100x in all four fragments and were included in further analyses. Of the excluded samples, two did not yield sequence in any fragment (Ct values 30.99 and 33.89) and three samples lacked complete sequence in one of four fragments (Ct values 20.58, 22.95, and 32.31). All five excluded samples were from 2016. Fig. 1 shows the coverage plot for a representative sample. The sequencing coverage for each sample is presented in supplementary table S3.

**FIG. 1.**
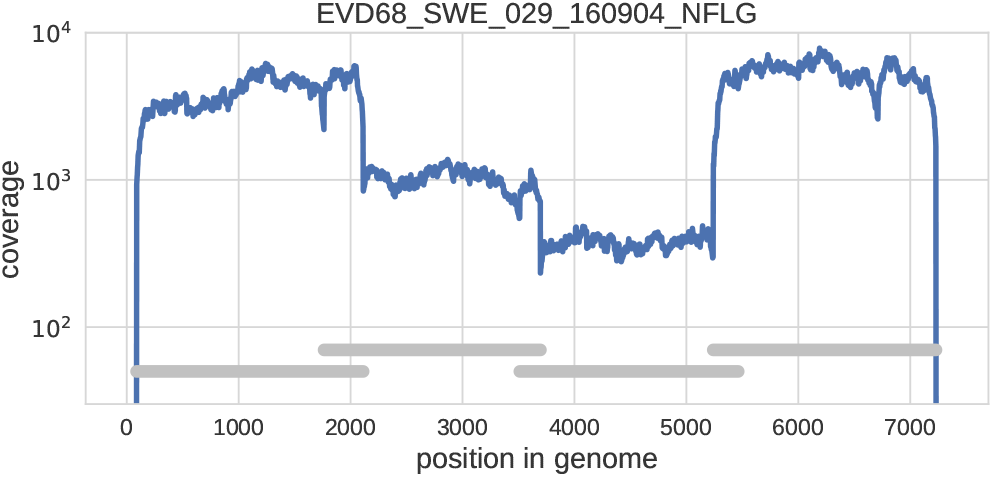
Example of coverage across the EV-D68 genome in a representative clinical sample. Note that the coverage varies somewhat in different parts of the genome due slightly different quantity of amplified DNA from the four overlapping amplicons in the material that was used for sequencing library preparation. The four overlapping are indicated by the horizontal grey bars.

### Reproducibility of identification of intrasample single nucleotide variants (iSNVs)

In addition to consensus sequences, deep population sequencing delivers information about intrasample diversity. Low genome template input, biased amplification, RT-PCR errors and sequencing errors can all potentially skew or inflate intrasample diversity estimate (Zanini *et al*., 2017). To control for such artifacts, we processed two high-titer patient samples in duplicate (see Methods). The accuracy of the detection and quantification of iSNVs was evaluated by comparing results from the two runs (Fig. 2A). Minor variants down to below 1% were consistently detected in both runs. Also, the estimated frequency of individual iSNVs showed good consistency between the two runs at frequencies above approximately 1% (Panel A), but a few iSNVs had markedly higher frequencies in one of the replicates (run 2 of sample SWE_037).

**FIG. 2.**
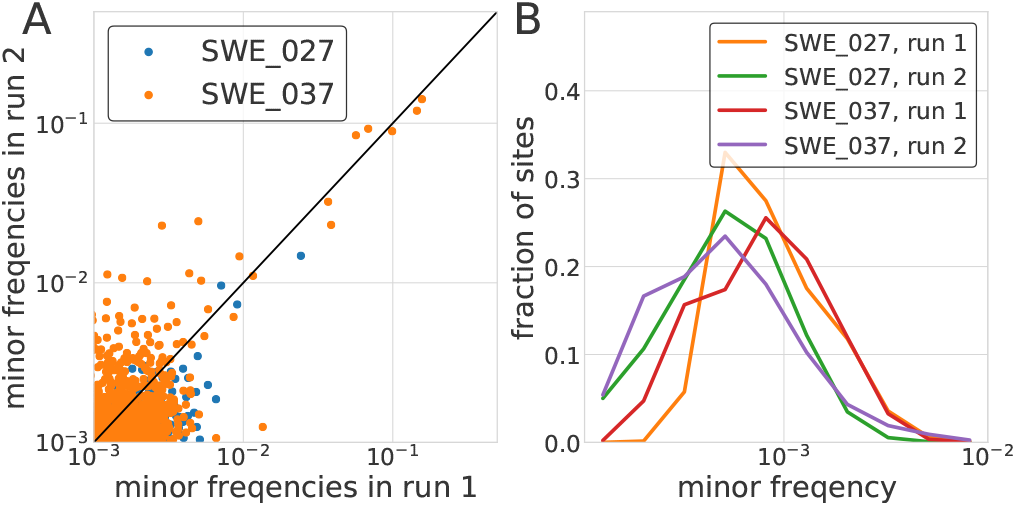
Accuracy of minor variant frequencies. Panel A shows the consistency of minor variant frequencies across two replicate extraction, RT-PCR, and sequencing of two samples. Minor variation was relatively consistently recovered down to a frequency of about 1%. Panel B shows the distribution of the frequencies of non-consensus calls across all sites covered in excess of 2, 000-fold. At the majority of sites, no variation above 0.001 is observed. Note that this variation includes within sample variation, RT-PCR errors, and sequencing errors.

At the majority of sites, we observed background of minor variation at frequencies well below 0.1% (see Fig. 2B).

Reproducibility of detection of iSNVs was also investigated in the five patients were sampled twice during their acute illness (0, 1, 1, 2 and 7 days apart). Supplementary figure S3 shows that frequencies of iSNVs above the 1% level were largely concordant across the two consecutive samples, especially in patient SWE_037 (for whom the first sample was also sequenced twice) and less so in patient SWE_012. In patient SWE_021, with seven days between the samples, there was a tendency for an increased variation in the second sample Unfortunately, the short time between the samples prevented us from estimating the rate of intrapatient evolution or other longitudinal aspects of intrapatient evolution of EV-D68.

Analysis of intrasample variability in all 54 patient samples also provided information about our ability to detect true biological variation. Fig. 3A shows the distribution of sites with variation above a certain frequency cutoff among the three codon positions. The distribution was approximately even between codon sites at very low cutoffs below 10^−3^ and then rapidly changed around approximately 0.3% before plateauing at 1%.

**FIG. 3.**
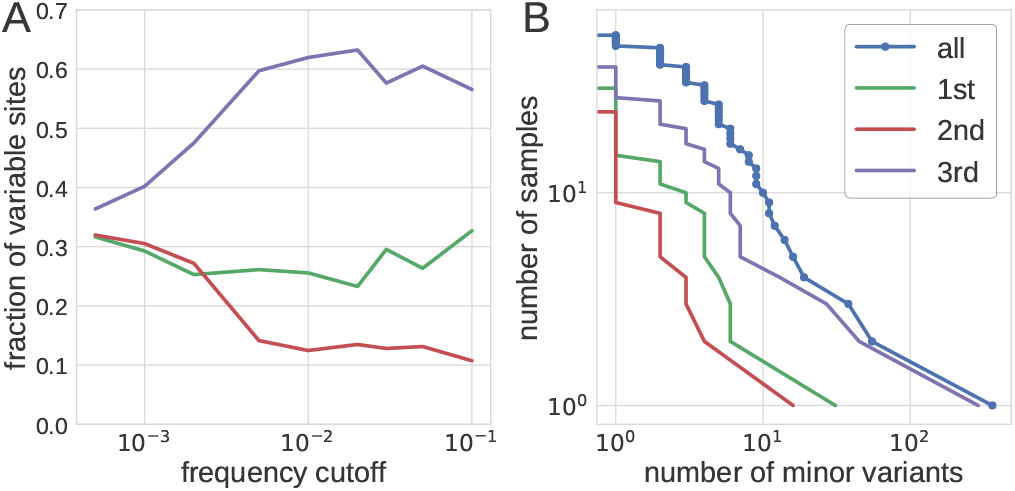
Within sample diversity. Panel A shows the distribution of the number of variable sites in coding regions among different coding positions. At very low frequency cutoffs, variable sites are approximately equally distributed in codons. The distribution rapidly changes once the frequency cutoff exceeds 0.3%, beyond which variable sites are found predominantly at 3rd codon positions and more rarely at 1st and 2nd codon positions. Panel B shows inverse cumulative distribution of the number of samples that have more than a certain number of sites that are variable above a level of 3%. Separate distributions are shown for all sites and for 1st, 2nd, and 3rd codon position in the coding region of the genome.

Collectively, our results show that a majority of iSNVs at frequencies >1% represent true biological variation, whereas variation <0.1% mostly is due to amplification and sequencing errors. Between 0.1% and 1% an increasing proportion of variation represents true iSNVs. The level of accuracy achieved is comparable to what we reported earlier for a similar sequencing strategy for HIV-1 (Zanini *et al*., 2017, 2016) and other viruses (Grubaugh *et al*., 2018).

### Intrapatient variability

As described above intrasample variability exceeding 1% likely represents true biological variation.

Approximately 60% of the variable sites above this level occurred at the 3rd codon position and around 30% and 10% in the 1st and 2nd codon positions, respectively (see Fig. 3A). Consequently, intrapatient variability is mostly synonymous (10%, 0%, and 95% of 1st, 2nd, and 3rd positions, respectively, admit synonymous mutations in our mapping reference).

Figure 3B shows that most (44 of 54) samples had fewer than 10 sites that displayed variability above a level of 3%. However, three samples had more than 20 variable sites, which suggested dual infections (see below). In the remaining 51 samples, both synonymous and non-synonymous substitutions were mostly scattered across the genome (Fig. 4). However, there was some clustering of minor variation in 1st and 2nd codon positions in the structural proteins (VP4 – VP1) including putative targets for neutralizing antibodies in BC and DE loops of VP1 (Liu *et al*., 2015). The details about complete amino acid substitutions and minor variation in these regions are given in supplementary tables S4 and S5.

**FIG. 4.**
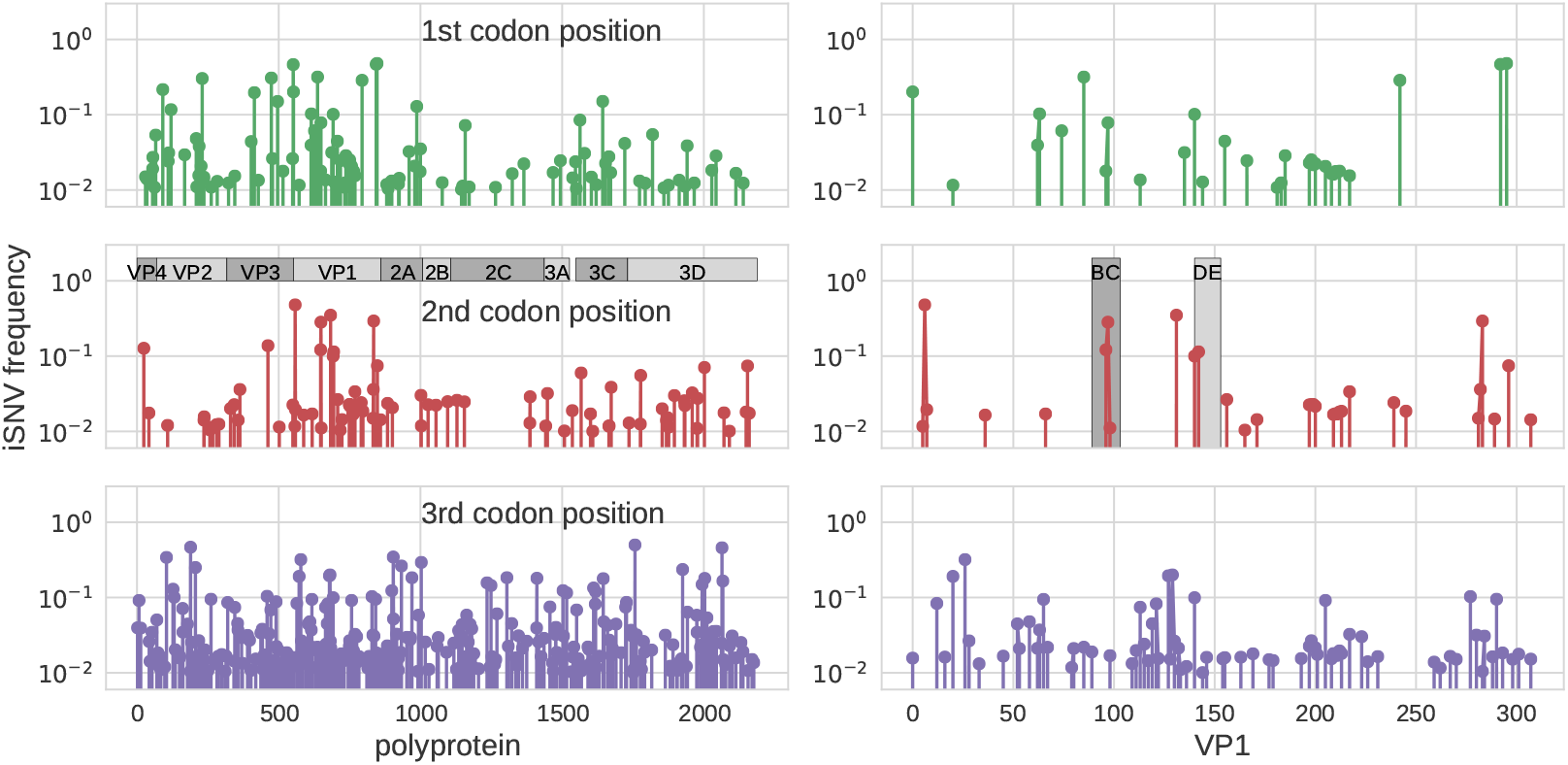
EV-D68 intrapatient variability across the genome and codon positions. The three rows show how intrapatient variability is distributed across the EV-D68 genome for 1st, 2nd, and 3rd codon position, respectively. The panel on the left show iSNVs along the entire polypeptide, the panels on the right zoom into VP1 and highlight the BC and DE loops. This figure includes data from one sample per patient (the first sample, for patients sampled twice), but excludes the three dual infected samples (see main text).

### Evidence of dual infections

As mentioned above, three samples (SWE_007, SWE_045, and SWE_046) drawn from three different patients showed more than 20 iSNVs. In all three samples, most minor variants were detected at a frequency around 10% (Fig. 5, left). This suggested the presence of two distinct variants; a major variant constituting around 90% of the virus population in the sample and a minor variant constituting around 10%. To investigate the origin of these minor variants we constructed a “minor” consensus sequence for each sample by exchanging the strict majority rule consensus nucleotide by the minor variant at positions where they exceeded a frequency of 1%. A phylogenetic tree analysis showed that the major and minor sequences were monophyletic in all samples except the three samples with indications of dual infections. For these samples the major and minor consensus sequences occupied different positions in the tree, see supplementary figure S1. To further investigate these putative dual infections we analyzed the linkage of iSNVs (Fig. 5, right). Almost all iSNVs occurred in complete linkage (LH=1) with neighboring iSNVs, which provides strong evidence for the presence of two distinct variants, rather than rapid evolution of intrapatient variation starting from a single transmitted variant.

**FIG. 5.**
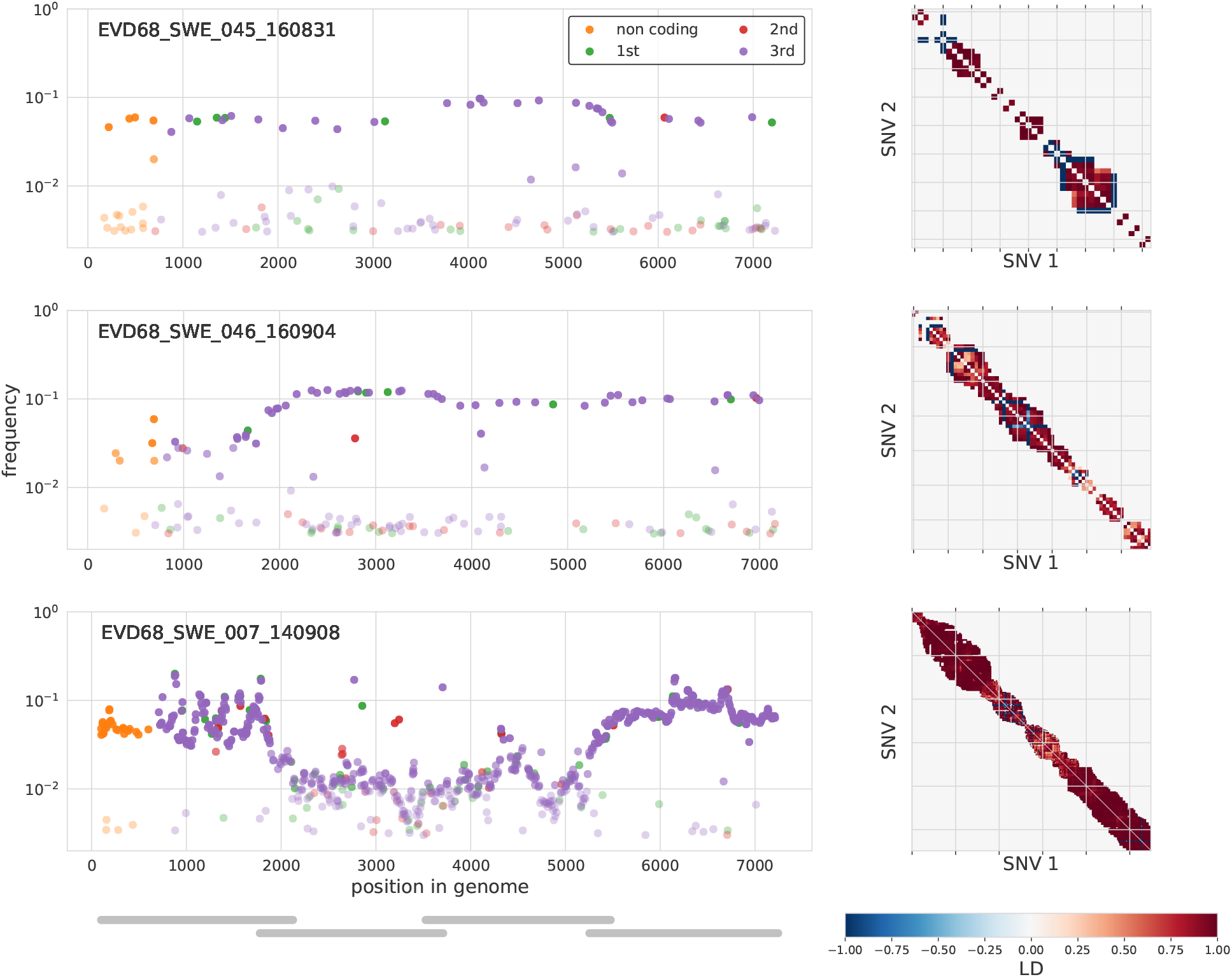
Minor iSNVs across the EV-D68 genome for three samples with putative dual infections. These three samples had more than 20 highly covered sites with minor variants in excess of 3% frequency. On the left, iSNVs are colored by codon position or as non-coding, and given opacity depending on their frequency. The majority of these variants were at 3rd positions and had similar frequencies across the amplicons, indicated by grey bars in the lower left panel. iSNVs in three amplicons (amplicon 1 of sample SWE_046 and amplicons 2 and 3 of sample SWE_007), however, were found at substantially lower frequency, possibly due to primer mismatches. The three panels on the right are indexed by iSNV order on the genome and show linkage disequilibrium between iSNVs (>1%) close enough to each other that they were covered at least 100-fold by the same sequencing read. Almost all of these variants at in complete linkage (dark red). A number of iSNVs just above 1% in sample SWE_045 and SWE_046 are likely variants in the background of the dominant variant. Those are in complete ‘anti-linkage’ with neighboring iSNVs are ~ 10% (dark blue).

### Phylodynamics of EV-D68

The interpatient evolution and phylogenetic relationship of the EV-D68 variants from Sweden and 509 published EV-D68 sequences was investigated using the `nextstrain` platform available at https://nextstrain.org/enterovirus/d68 (Hadfield *et al*., 2018), which allows users to interactively explore the dataset. A screen shot is displayed in Fig. 6.

**FIG. 6.**
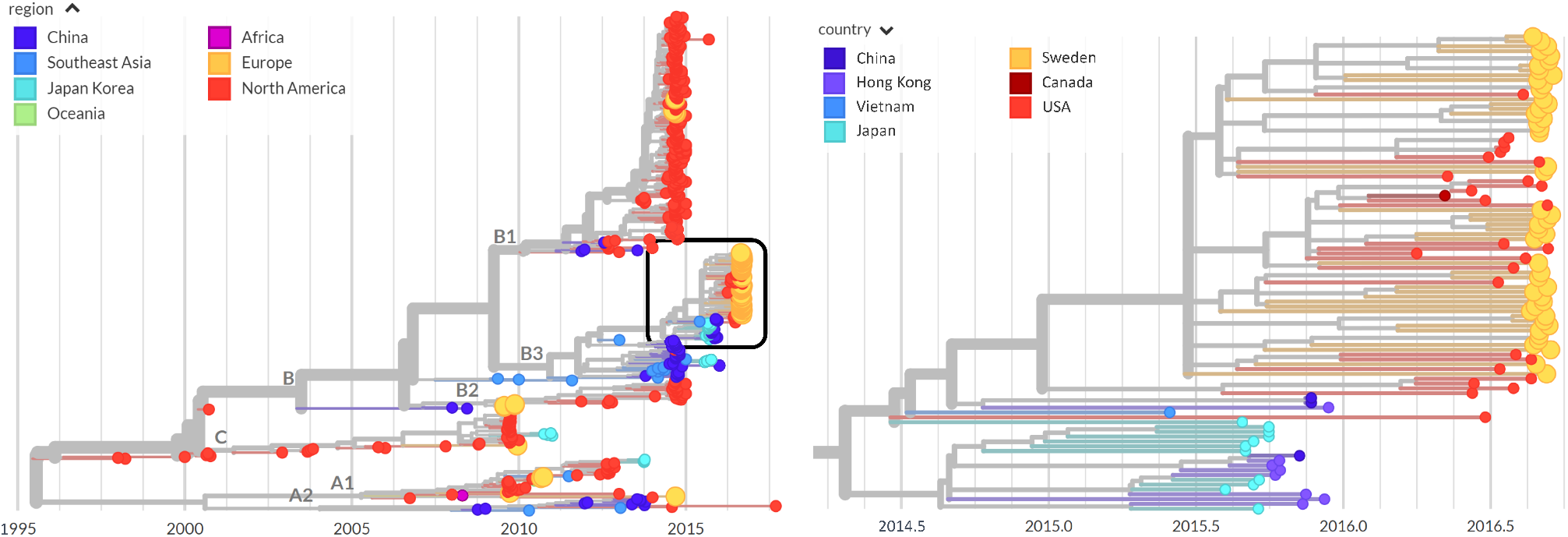
Phylogenetic analysis. The left panel shows a phylogenetic tree as rendered by `nextstrain` with the sequences labelled by region. Those from Europe, including those from Sweden, are colored in orange. The right panel is a zoomed-in view of the boxed area on the left-hand panel colored by country and highlighting the Swedish sequences from the 2016 outbreak (orange). Swedish sequences often cluster together, but are interspersed with sequences from Canada (dark red) and the USA (red), implying multiple introductions that then spread locally.

The `nextstrain` phylodynamic analysis showed that all Swedish EV-D68 variants from the 2016 outbreak belonged to the B3 lineage of EV-D68, which agrees with our earlier findings based on VP2/VP4 gene sequences (Dyrdak *et al*., 2016). Within the B3 lineage our new Swedish sequences were interspersed with international sequences, primarily from the US, but monophyletic clusters of Swedish sequences were also observed (Fig. 6, right). A bootstrap analysis showed that most branches were well supported (Fig. S4).

These findings indicate that during the worldwide 2016 outbreak EV-D68 was introduced into Stockholm several times, and that these separately introduced variants thereafter spread locally.

Interestingly, the tree indicated that the 2016 B3 sub-clade did not evolve directly from the 2014 B1 subclade. Instead, the most recent common ancestor (MRCA) of the B1 and B3 subclades was estimated to have existed in first half of 2009 (90% CI Dec 2008 – Jul 2009). Many of the proximal sequences in the B1 and B3 subclades were sampled in East Asian countries, but this should be interpreted with caution since sampling probably has been incomplete and geographically biased. A tangle-tree analysis suggested recombination within subclades, but not between clades or subclades (supplementary figure S2), but also this finding should be interpreted with caution since there was limited intrasubclade variation.

### Interpatient evolution

A plot of root-to-tip distances of EV-D68 sequences vs. time showed an extremely strong temporal signal (Fig. 7). We estimated an evolutionary rate of 3.8 · 10^−3^ ± 10^−4^ substitutions per site and year for the entire genome and 4.2 · 10^−3^ ± 4 · 10^−4^ for the VP1 region, which is slightly lower than previous estimates (Ny *et al*., 2017; Tokarz *et al*., 2012)). 87% of all substitutions were synonymous.

**FIG. 7.**
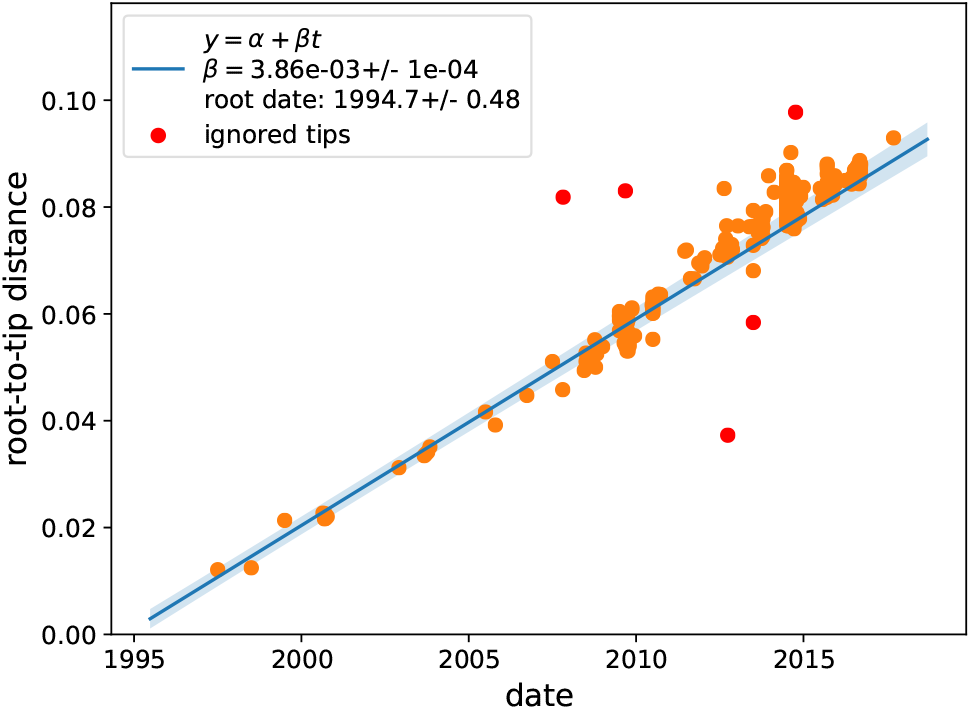
Temporal signal. A scatter plot of root-to-tip distance vs time indicates an extremely strong temporal signal with an evolutionary rate of *μ* = 0.0039± 0.0001 substitutions per site and year.

We used `nextstrain` to investigate diversity across the EV-D68 genome in the sample containing the genomes reported in this study and database genomes. Ancestral sequences were reconstructed using the phylogeny in Fig. 6, and the number of transition events for each codon were counted along the tree. Fig. 8 shows that amino acid substitution events were observed in all EV-D68 proteins and varied between 0 and 27 events per codon. The five positions with the highest number of substitution events were: codon 234 in protein VP3 (27 events), 273 in 2C (20 events), 22 in 2A (15 events), 60 in VP3 (15 events), and 277 in 2C (14 events). None of these positions have been reported to be neutralizing epitopes. Codon 277 in 2C and codon 1 in VP1 showed evidence of episodic diversifying selection as per MEME analysis (*p* < 0.05).

**FIG. 8.**
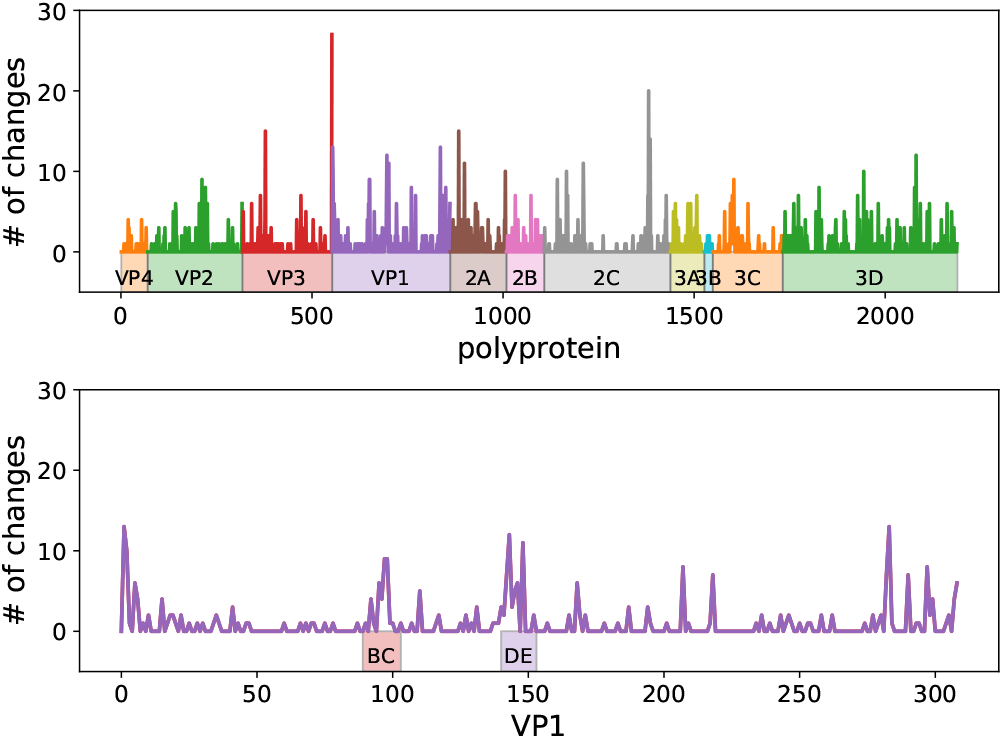
Interpatient diversity of EV-D68. Amino acid substitution events on the phylogeny in Fig. 6 left were enumerated using `nextstrain`. Top panel shows the number of such events for each codon along the entire coding sequence. Bottom panel shows events in VP1, where peaks were observed in the BC and DE loops, as well as both ends of the protein.

The BC and DE loops in the VP1 protein showed relatively high diversity, especially in positions 95, 97, 98, 142, 143 and 148, suggesting that immune selection may have impacted on EV-D68 evolution (see Fig. 8, bottom panel, and for details see `nextstrain`).

We also used `nextstrain` to track on which branches in the EV-D68 phylogeny amino acid substitutions had evolved and been fixated. The branches leading to the MRCA of the B1 and B3 clades had many amino acid substitutions, including substitutions at codons 90, 95, 97, 98, 145 and 148 in the BC and DE loops. In contrast, no amino acid substitutions in the loops were observed on branches separating the B1 and B3 subclades. Similarly, only very few substitutions in the loops were observed within the B1 and B3 subclades.

## DISCUSSION

We have performed a comprehensive investigation of the intra- and interpatient evolution of EV-D68 using new deep near full-length sequences from 54 Swedish patients sampled during the 2014 and 2016 outbreaks and available database sequences. The new data and previously published sequences are available on `nextstrain` to facilitate real-time tracking and further study of EV-D68 evolution and spread. Intrapatient variability was mostly low and dominated by rare synonymous substitutions, consistent with deep sequencing studies on in fluenza virus infections (Debbink *et al*., 2017). However, three patients showed evidence of dual infections. The phylodynamic analysis indicated EV-D68 was introduced into Stockholm several times during the 2016 outbreak and then spread locally.

Three patients showed evidence of dual infection with two different EV-D68 variants. Even though co-infections with different enteroviruses has been reported for poliovirus (PV) and non-polio enteroviruses (NPEV) (Isaacs *et al*., 2018; Melnick *et al*., 1951; Parks *et al*., 1967), this is to our knowledge the first report of dual infections with EV-D68. Dual infections during the 2016 outbreak are not completely surprising in view of the high EV-D68 incidence during the outbreak (Dyrdak *et al*., 2016) and the fact that the present study showed multiple introductions of the B3 subclade into Stockholm. We also found indications of intrasubclade recombination, but not between clades or subclades. Similar findings have been reported in three recent studies (Ny *et al*., 2017; Tan *et al*., 2015; Yip *et al*., 2017). It is expected that recombination primarily would occur within sub-clades since recombination requires co-infection with two or more virus variants and EV-D68 variants that circulate during an outbreak or season tend to be closely related.

The EV-D68 genome was amplified in four overlapping amplicons that were sequenced on the Illumina HiSeq platform. Our results indicate that variation down to a frequency of 1% represented “true” iSNVs, which were reproducibly detected and quantified. In contrast, most variation below 0.1% represented noise from cDNA synthesis, PCR and sequencing. It would be interesting to compare the performance of our sequencing protocol with the pan-enterovirus protocol recently published by (Isaacs *et al*., 2018). In the 54 samples that were sequenced to sufficient depth we observed that most nucleotides were conserved. Among iSNVs detected above the 1% level, most occurred at the 3rd codon position, i.e. were usually synonymous. The finding that intrapatient variation was limited and mostly synonymous was expected since infection usually appeared to have been established by a single virion and EV-D68 is an acute infection where is limited time for the virus to diversify within the infected host.

Albeit there is a bias introduced by sampling and typing being done only at certain centras, the phylodynamic analysis showed that the splits between EV-D68 clades and subclades occurred several years ago. For instance, the MRCA of the B1 and B3 subclades was estimated to have existed in 2009. Enterovirus infections, like influenza, has a seasonal pattern. However, in contrast to influenza, which infects people of all age groups and has a ladder-like phylogenetic pattern that indicates a strong immune selection, EV-D68 infects predominantly children, and its phylogenetic pattern does not suggest continuous immune escape (Grenfell *et al*., 2004). In line with this, the branches leading to the B1 and B3 sub-clades did not have amino acid substitutions in the BC and DC loops, which are putative targets for neutralizing antibodies. However, it should be acknowledged that there is limited knowledge about targets for humoral and cellular immunity in EV-D68.

Our phylodynamic analysis suggested that the recent global outbreaks of EV-D68 might have been preceded by low-level circulation of EV-D68 in East Asia. However, this pattern should be interpreted with much caution since the number of available full-length EV-D68 sequences is low and the sequences have not been systematically collected in time or space. We hope that the `nextstrain` platform will allow for continuous collection and sharing of EV-D68 sequence data, which will be needed to obtain a better understanding of how new successful EV-D68 variants evolve and spread. Better knowledge about humoral and cellular immunity against EV-D68 is also needed.

## ACKNOWLEDGEMENTS

We gratefully acknowledge expert sequencing service and advice by Karin Sollander, Cecilia Svensson and Valtteri Wirta at the Clinical Genomics Unit at Science for Life Laboratory (SciLifeLab), and Lina Thebo and Eva Eriksson at the Karolinska Institute. We would also like to thank the entire `nextstrain` team.

## SUPPLEMENTARY INFORMATION

**TABLE S1.**
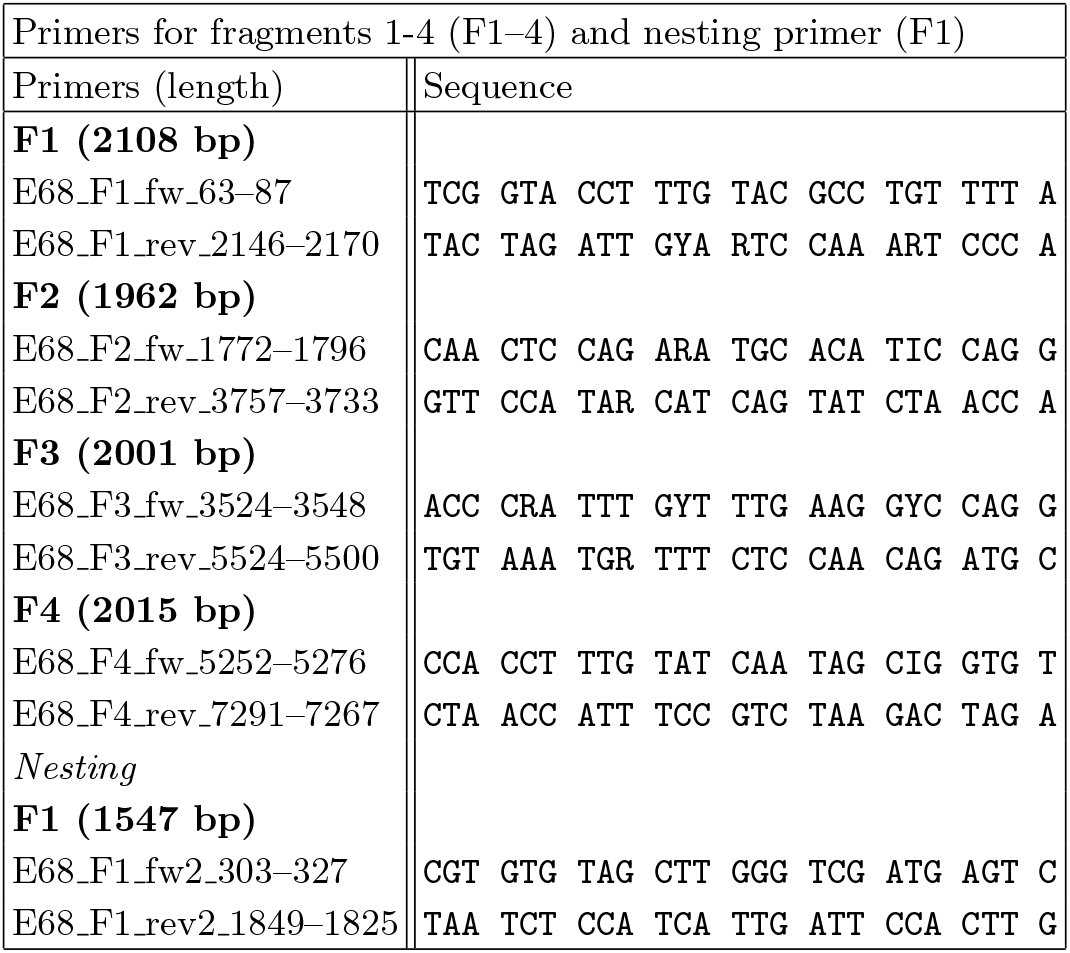
Primers for near full-length sequencing of EV-D68 in four overlapping fragments (F1–F4). The nested primers for F1 were used for template quantification.

**TABLE S2.**
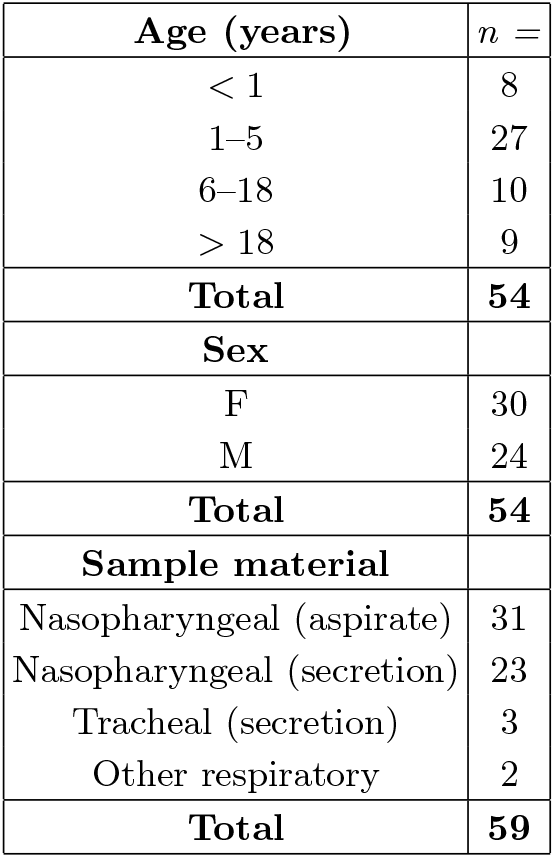
Breakdown of patient age, sex, and sample material.

**TABLE S3.**
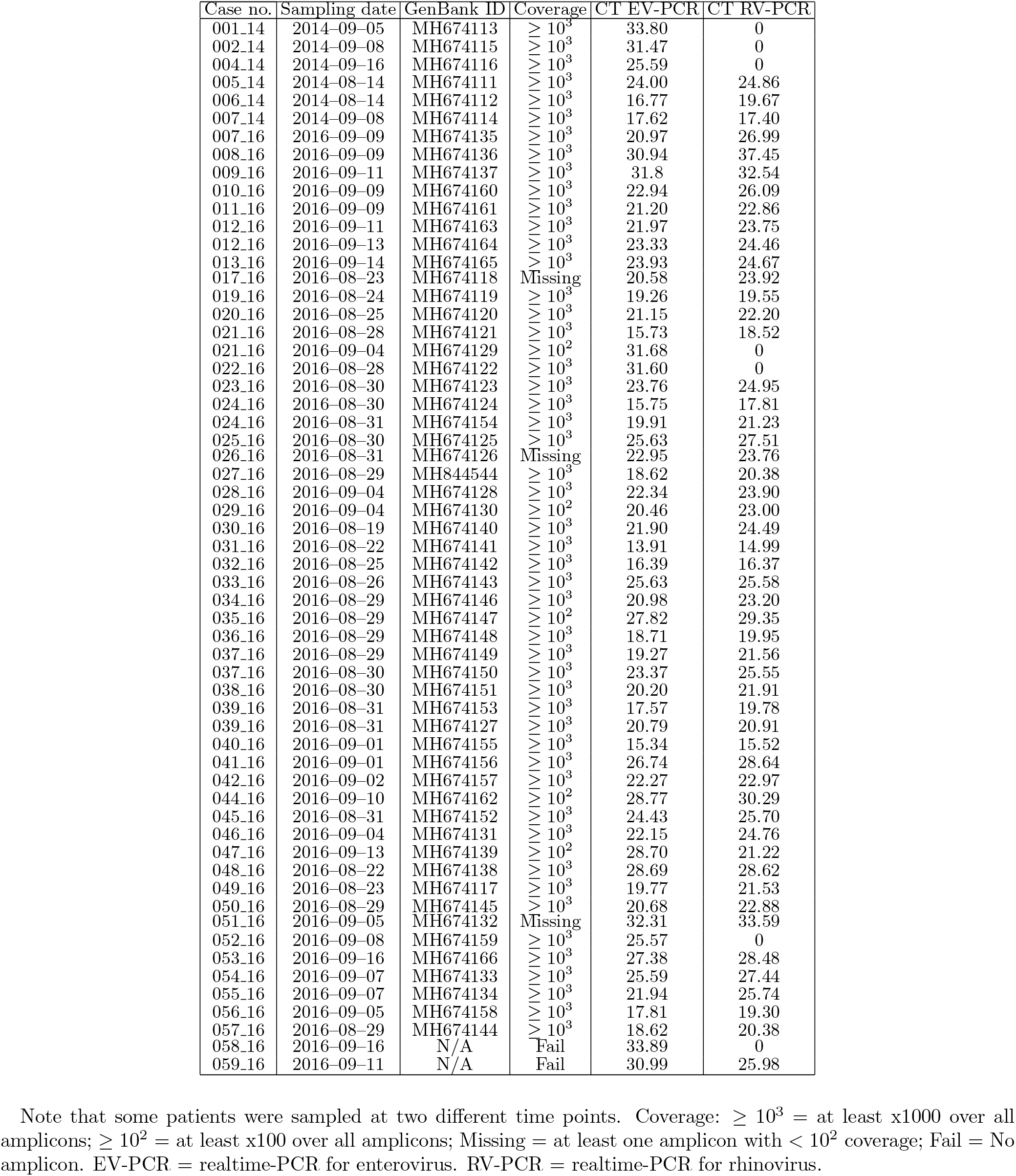
Details about study samples.

**TABLE S4.** Amino acid sequence and substitutions in the putative targets for neutralizing antibodies in the BC-loop in the VP1 protein (codon 89–105, consensus sequence KDHTSSAAQADKNFFKW).

| Sample |  |  | Codon position and consensus amino acid |  |  |  |
| --- | --- | --- | --- | --- | --- | --- |
| Case no. | Collection date | Sequence variant | 96<br>A | 98<br>Q | 99<br>A | 100<br>D |
| 007_16 | 2016-09-09 | Major | - | - | T | - |
| 007_16 | 2016-09-09 | Minor | - | - | T | - |
| 012_16 | 2016-09-11 | Major | - | - | T | - |
| 012_16 | 2016-09-11 | Minor | - | - | T | - |
| 012_16 | 2016-09-13 | Major | - | - | T | - |
| 012_16 | 2016-09-13 | Minor | - | - | T | - |
| 021_16 | 2016-09-04 | Minor | - | R | - | - |
| 022_16 | 2016-08-28 | Major | - | - | T | - |
| 022_16 | 2016-08-28 | Minor | - | - | T | - |
| 023_16 | 2016-08-30 | Major | - | - | T | - |
| 025_16 | 2016-08-30 | Minor | - | - | - | H |
| 027_16 | 2016-09-01 | Major | - | - | T | - |
| 027_16 | 2016-09-01 | Minor | - | - | T | - |
| 028_16 | 2016-09-04 | Minor | - | R | - | - |
| 031_16 | 2016-08-22 | Major | - | - | T | - |
| 031_16 | 2016-08-22 | Minor | - | - | T | - |
| 033_16 | 2016-08-26 | Minor | - | - | - | H |
| 036_16 | 2016-08-29 | Major | - | - | T | - |
| 036_16 | 2016-08-29 | Minor | - | - | T | R |
| 038_16 | 2016-08-30 | Minor | - | - | V | - |
| 039_16 | 2016-08-31 | Minor | - | R | - | - |
| 053_16 | 2016-09-16 | Major | - | - | T | - |
| 053_16 | 2016-09-16 | Minor | - | - | T | - |
| 054_16 | 2016-09-07 | Major | T | - | - | - |
| 054_16 | 2016-09-07 | Minor | T | - | - | - |

**TABLE S5.**
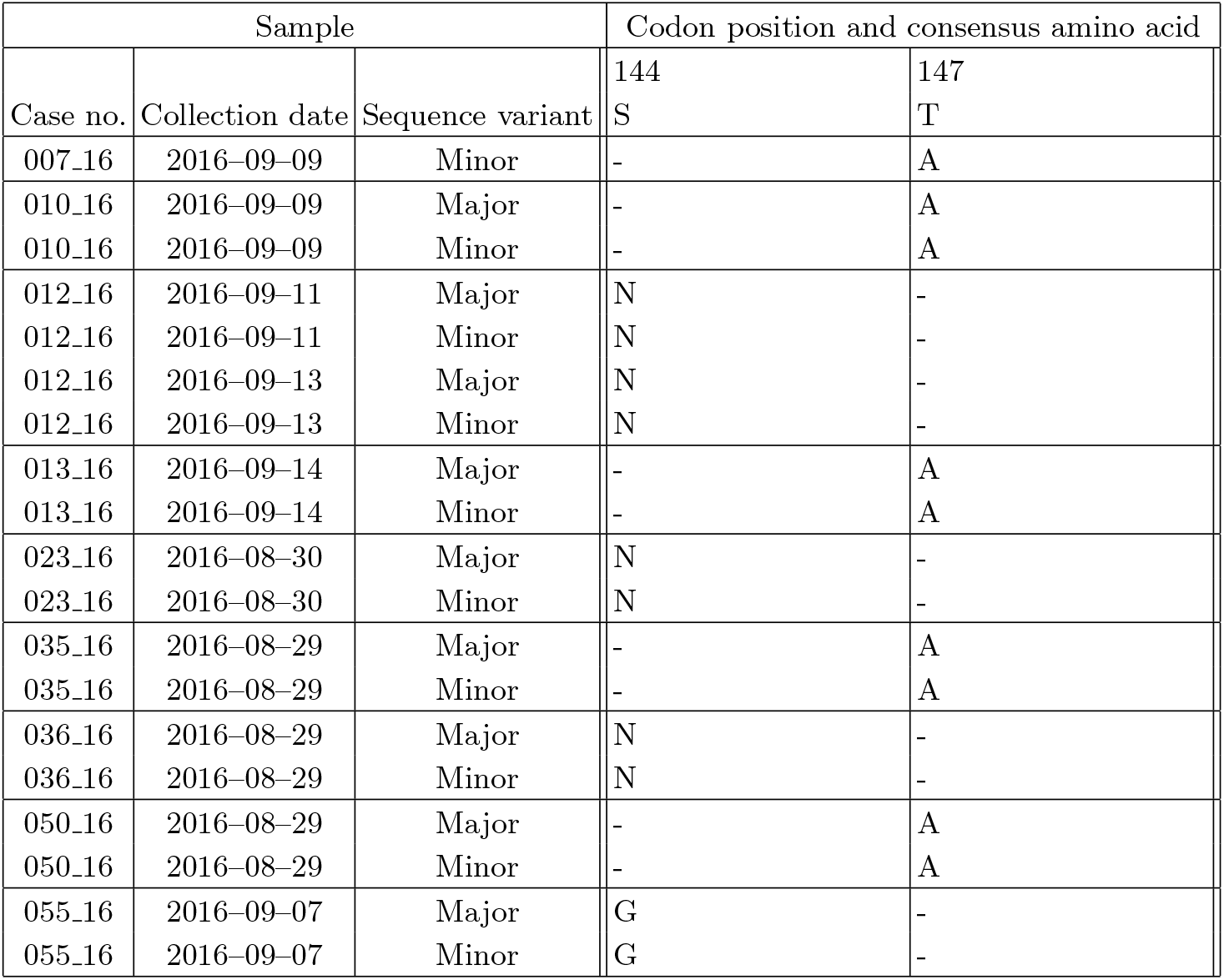
Amino acid sequence and substitutions in the putative targets for neutralizing antibodies in the DE-loops in the VP1 protein (codons 140–152 consensus sequence: NGSSNNTYVGLPD).

| Sample |  |  | Codon position and consensus amino acid |  |
| --- | --- | --- | --- | --- |
| Case no. | Collection date | Sequence variant | 144<br>S | 147<br>T |
| 007_16 | 2016–09–09 | Minor | - | A |
| 010_16 | 2016–09–09 | Major | - | A |
| 010_16 | 2016–09–09 | Minor | - | A |
| 012_16 | 2016–09–11 | Major | N | - |
| 012_16 | 2016–09–11 | Minor | N | - |
| 012_16 | 2016–09–13 | Major | N | - |
| 012_16 | 2016–09–13 | Minor | N | - |
| 013_16 | 2016–09–14 | Major | - | A |
| 013_16 | 2016–09–14 | Minor | - | A |
| 023_16 | 2016–08–30 | Major | N | - |
| 023_16 | 2016–08–30 | Minor | N | - |
| 035_16 | 2016–08–29 | Major | - | A |
| 035_16 | 2016–08–29 | Minor | - | A |
| 036_16 | 2016–08–29 | Major | N | - |
| 036_16 | 2016–08–29 | Minor | N | - |
| 050_16 | 2016–08–29 | Major | - | A |
| 050_16 | 2016–08–29 | Minor | - | A |
| 055_16 | 2016–09–07 | Major | G | - |
| 055_16 | 2016–09–07 | Minor | G | - |

**FIG. S1.**
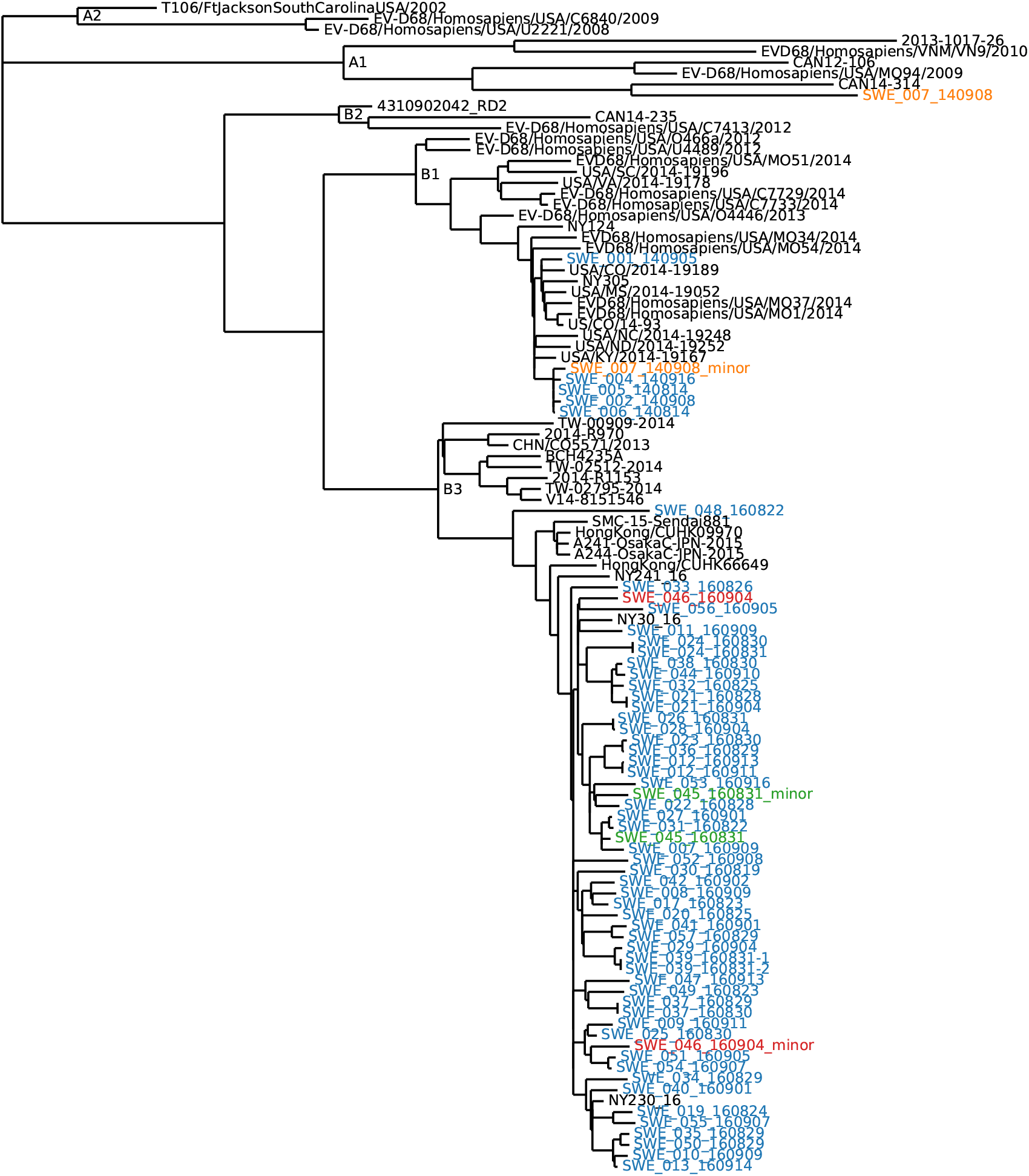
Phylogenetic placing of minor variants in dually infected patients. The figures hows a mid-point rooted ML tree of all consensus sequences obtained in this study (in blue) along with minority sequences from three samples from dually infected individuals (in orange, red, and green), and a sample of publicly available sequences to illustrate some EVD68 clades (labelled on internal nodes). The minor variants do not cluster tightly with any other sample but rather look like representative samples from the populations. Minor variant sequence of SWE_007_140908 is slightly problematic since variants in the amplicons F2 and F3 cannot be cleanly separated from sequencing error. To control for this, the region from 2,000-5,500bp in this sequence was replaced with ’N’s.

**FIG. S2.**
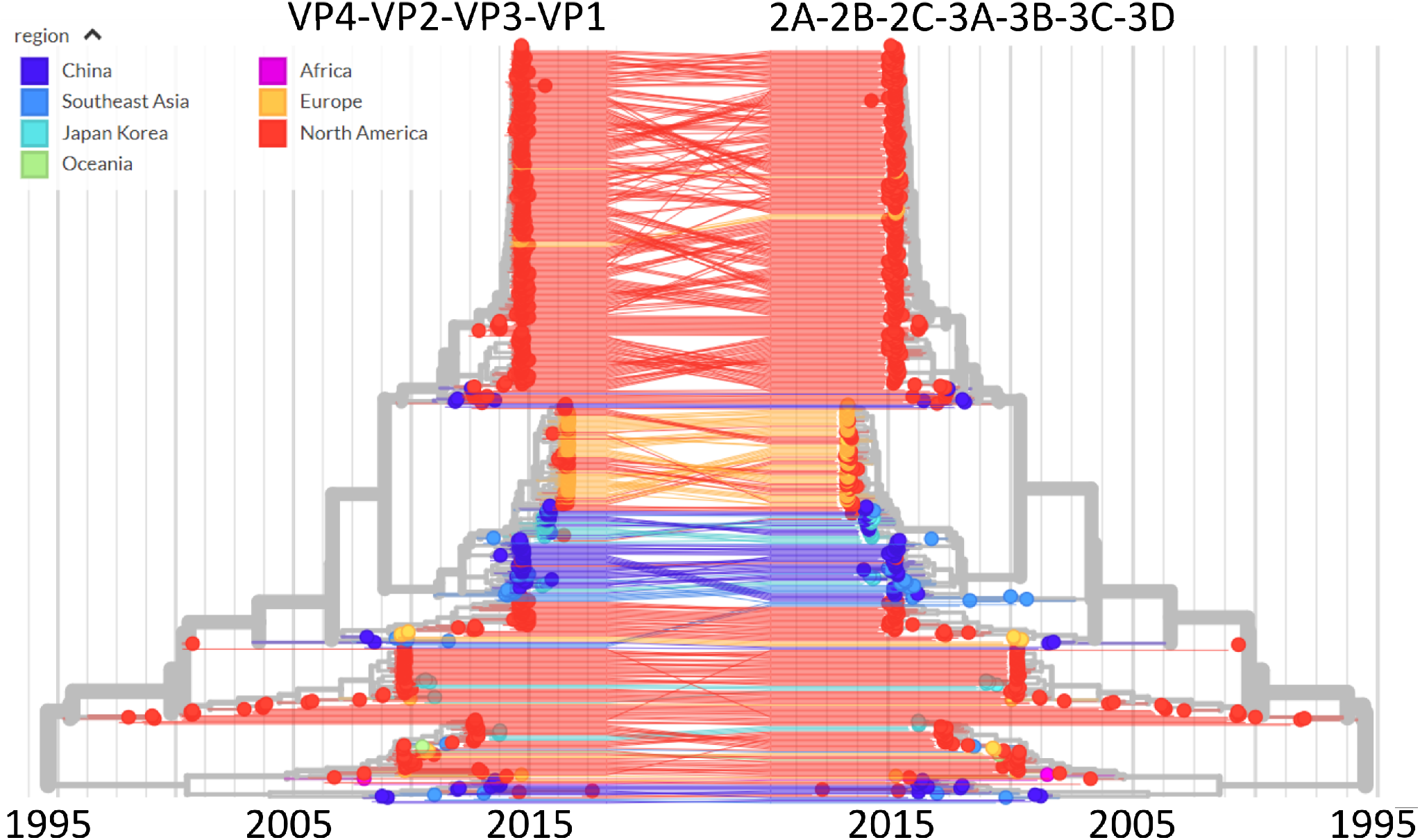
Tangle-tree analysis for possible EV-D68 recombinants. Phylogenetic trees of sequences coding for VP4-VP1 (5’ region) and 2A-3D (3’ region) are congruent at the level of major clades, suggesting little inter-clade recombination. Trees were calculated using the nextstrain analysis pipeline and visualized using nextstrain’s auspice.

**FIG. S3.**
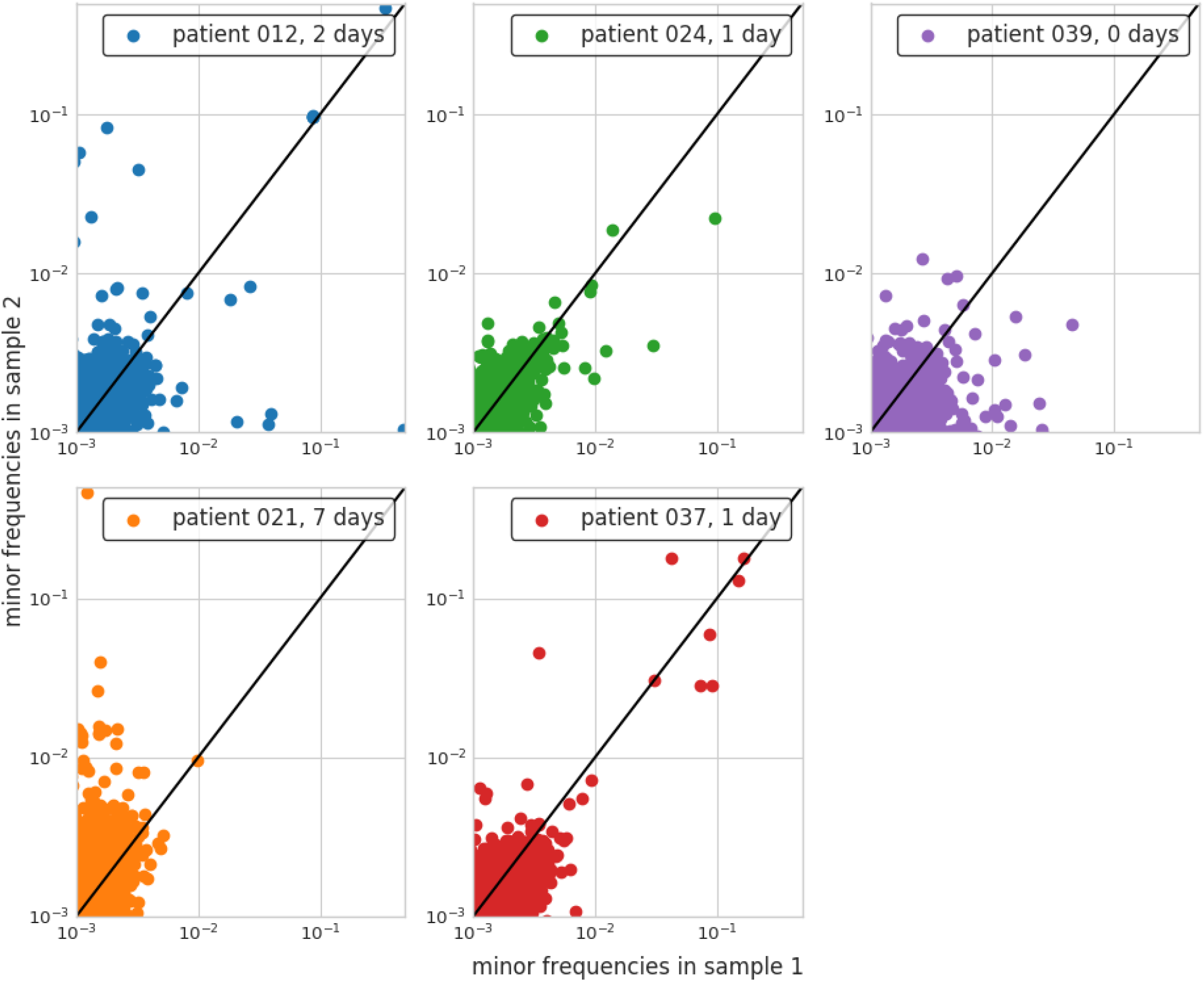
Minor variant consistency in repeatedly sampled individuals. Each panel shows a scatter plot of minor variant frequencies in two samples from the same individual, sampled between 0 and 7 days apart. As expected, most iSNVs are at frequencies below one percent but some samples harbor iSNVs in the range of 10%. Their frequencies are largely concordant for samples 1 day apart, but deviate in the samples that are 2 or 7 days apart.

**FIG. S4.**
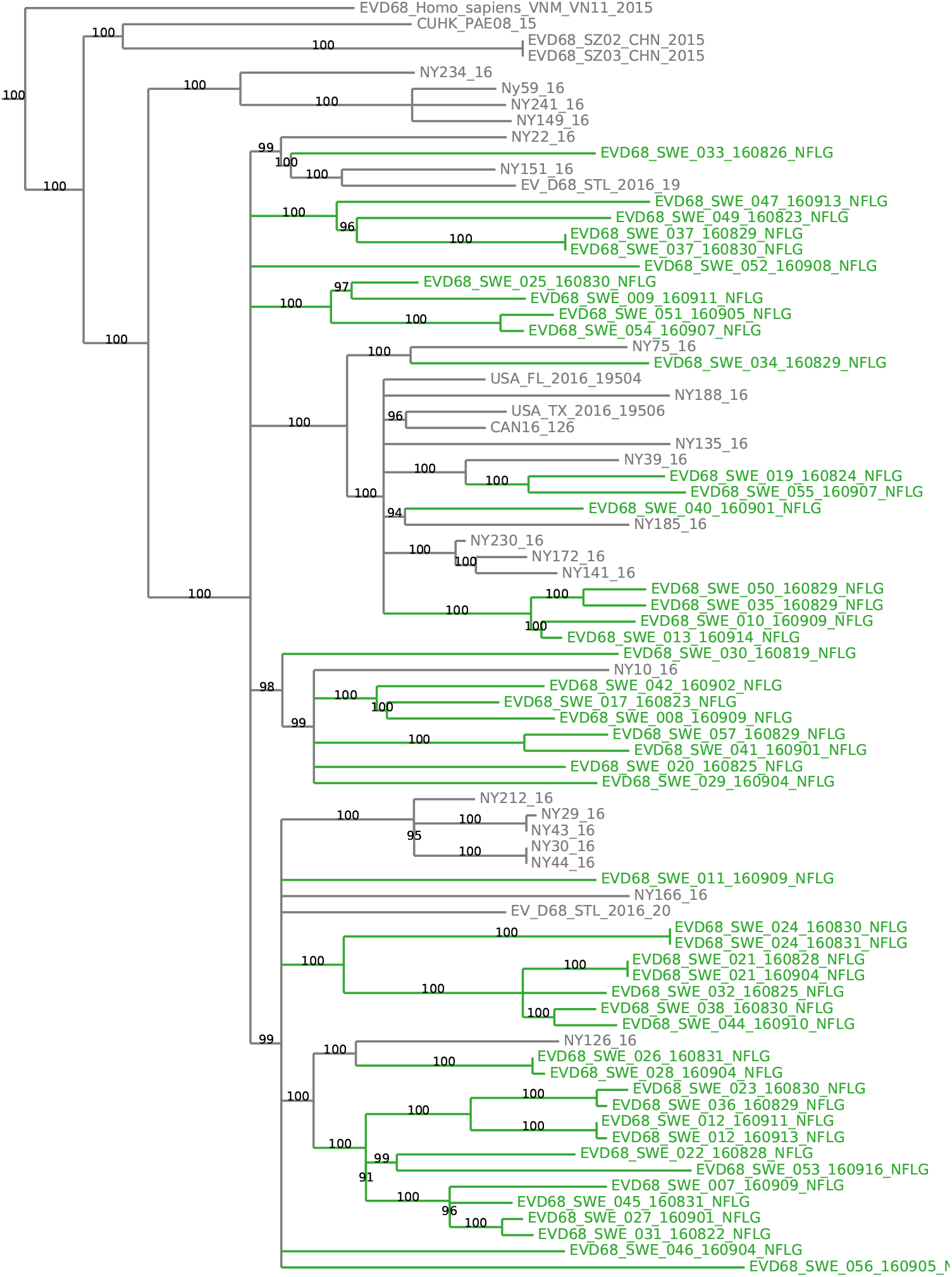
Swedish samples are not monophyletic. This figures show a tree in which all branches with less than 90% bootstrap support are collapsed (bootstrap values are shown on remaining branches) and monophyletic clusters of exclusively Swedish samples are colored in green. Even after reducing the tree to well supported branches, Swedish samples are interspersed with North American samples. There are at least eight “Sweden only”-clades that are separated from other Swedish clades by international isolates.

